# Candidate genes under balancing selection in a plant bacterial pathogen

**DOI:** 10.1101/388207

**Authors:** José A. Castillo, Spiros N. Agathos

## Abstract

Plant pathogens are under significant selective pressure by the plant host. Consequently, they are expected to have adapted to this condition or contribute to evading plant defenses. In order to acquire long-term fitness, plant bacterial pathogens are usually forced to maintain advantageous genetic diversity in populations. This strategy ensures that different alleles in the pathogen’s gene pool are maintained in a population at frequencies larger than expected under neutral evolution. This selective process, known as balancing selection, is the subject of this work in the context of a common plant bacterial pathogen. We performed a genome-wide scan of *Ralstonia solanacearum,* an aggressive plant bacterial pathogen that shows broad host range and causes a devastating disease called ‘bacterial wilt’. Using a sliding window approach, we analyzed 57 genomes from three phylotypes of *R. solanacearum* to detect signatures of balancing selection. A total of 161 windows showed extreme values in three summary statistics of population genetics: Tajima’s D, Watterson’s θ and Fu & Li’s D*. We discarded any confounding effects due to demographic events by means of coalescent simulations of genetic data. The prospective windows correspond to 78 genes that map in any of the two main replicons of *R. solanacearum.* The candidate genes under balancing selection are related to primary metabolism (51.3%) or directly associated to virulence (48.7%), being involved in key functions targeted to dismantle plant defenses or to participate in critical stages in the pathogenic process. These genes are useful to understand and monitor the evolution of bacterial pathogen populations and emerge as potential candidates for future treatments to induce specific plant immune responses.

## INTRODUCTION

Balancing selection (BS) is a well-known concept in evolutionary biology and population genetics that has extensively analyzed in many organisms. BS is a type of positive selection that favors the maintenance of a high genetic diversity within a given population. This diversity could be displayed as an excess of polymorphisms on existing alleles or as the maintenance of different alleles at selected loci. Usually BS influences genetic variation in genomes in a localized way, maintaining diversity at the selected sites but also increasing diversity at closely linked neutral sites (7). BS works through different mechanisms, namely, heterozygote advantage (also called overdominant selection) (26), frequency-dependent selection (58) and spatial/temporal heterogeneity (27). One particularly interesting case is frequency-dependence selection that is related to the coevolution between host and pathogen following the ‘trench warfare’ model. This model postulates that coevolution of both host and pathogen leads to stable richness of polymorphisms through BS (59). Good examples of this model are interactions of plant resistance genes with virulence-related genes of the pathogen under defined ecological and epidemiological conditions specific for each host-pathogen system. In this case, elevated polymorphism levels in pathogen virulence genes have been found in several systems and therefore it is possible to detect them using standardized tests (59).

Lately, much attention has been paid to BS in different eukaryotic species such as humans (33; 4), plants (52) and parasites (62), however, very little to bacteria, with one exception, two species of *Staphylococcus* genus that cause serious diseases in the respiratory tract, skin and other organs of humans (64; 71). At the level of plant bacterial pathogens, there are no reported works that indicate whether BS is a significant force that shapes populations, modulates the interaction with the plant host and directs evolution. In this work, we focus on detecting BS events in the far less studied plant host-bacterial pathogen system and take *Ralstonia solanacearum* as model species to perform the analyses.

*R. solanacearum* belongs to the Betaproteobacteria class and the Burkholderiaceae family and is considered a species complex (RSSC) because it is composed of a large number of genetic groups, often subdivisible into a number of clonal lines (22). RSSC has lately been re-classified in three different species (54) based on a previous phylogenetic arrangement that divided the complex into four phylotypes (15). Each phylotype constitutes a major monophyletic cluster of strains and reflects a geographic origin: phylotype I (Asia), phylotype II (Americas), phylotype III (Africa), and phylotype IV (Indonesia)(15; 5). Phylogenetic studies show that phylotype II is also divided into two monophyletic subgroups designated IIA and IIB (5). Strains belonging to this RSSC are aggressive plant pathogens that cause wilt disease of more than 250 plant species including economically valuable crops. These bacteria alternate between two lifestyles, as saprophytic on soil and water, and as pathogen inside plant tissues and organs. The bacteria enter susceptible plants through the roots, invade the xylem vessels, form biofilms and spread to the aerial parts of the plants. For pathogenesis, RSSC strains use an ample repertoire of molecular weapons like cell wall degrading enzymes, an extracellular polysaccharide and effectors secreted through the type III secretion system (T3SS) (20). All virulence factors are expressed and eventually secreted in a coordinated manner and appear to have additive effects since no single factor can completely explain infection and disease symptoms (38). At the genomic level, the RSSC strains harbor two DNA circular molecules, a larger replicon of 3.7 Mb and a smaller 2.1 Mb replicon, corresponding to chromosome and megaplasmid respectively. Both replicons contain housekeeping as well as virulence-related genes (55).

To investigate BS in the RSSC, we performed a genome-wide scan on both replicons (chromosome and megaplasmid) and attempted to determine whether BS is more frequent in essential versus virulence-related genes. Only for the purposes of this work, we have considered each RSSC phylotype (including subgroups IIA and IIB) as single, independent populations and have measured the excess of common polymorphisms using the classical summary statistics (Tajima’s D and others) rather than rely on model-based methods (13) or new summary statistics (like β)(56) because it was considered that they would not add more confidence to the results when used together with Tajima’s D.

## DATA AND METHODS

### Sequence data and alignment

Fifty-seven full-genome sequences of three RSSC phylotypes were downloaded from NCBI’s FTP server (https://www.ncbi.nlm.nih.gov/genome/microbes/) in February and April, 2017. We selected 20 genomes for phylotype I (CQPS-1, FJAT-1458, FJAT-91, FQY_4, GMI1000, KACC10709, OE1-1, PSS1308, PSS190, PSS4, RD13-01, RD15, Rs-09-161, Rs-10-244, Rs-T02, SD54, SEPPX05, TO10, UW757, YC45) and phylotype IIB (23-10BR, CFBP1416, CFBP3858, CFBP6783, CFBP7014, CIP417, GEO_304, GEO_96, IBSBF1503, IPO1609, Po82, RS 488, RS2, UW163, UW179, UW24, UW365, UW491, UW551, UY031). For phylotypes IIA and IV we used the largest number of genomes available in the database (12 genome sequences: B50, BBAC-C1, CFBP2957, CIP120, Grenada 9-1, IBSBF1900, K60, P597, RS 489, UW181, UW25, UW700; and 5 genome sequences: A2-HR MARDI, KACC 10722, PSI07, R229, R24, respectively). Unfortunately, there was not enough genome sequences for phylotype III at the time we retrieved sequences to perform analyses, therefore we did not include this phylotype in the analysis. All analyses were performed on the main (chromosome) and the secondary (megaplasmid) replicons of the RSSC.

We aligned the genome sequences using progressiveMauve aligner v2.4.0 (12) with default settings. For phylotype IV sequences, we increased the gap penalty (gap open score −600) to avoid opening unnecessarily large gaps, however we allowed small gaps (3-10 bp). For all analyses we used only Locally Collinear Blocks (LCBs) sequences to assure we worked with homologous sites that show maximal collinearity in order to avoid problems of internal genome rearrangements and gene gain and loss. We used stripSubsetLCBs script distributed with Mauve to extract LCBs longer than 1000 bp that were shared by RSSC genomes. This script generates an xmfa file that should be converted to a fasta file to facilitate the ensuing analyzes. For this purpose, we used a Perl script (xmfa2fasta).

### Statistical analyses

We applied summary statistics to detect BS. The summary statistics were used to measure an excess of polymorphisms linked to the genomic regions under this type of selection. We adopted three different statistics: Watterson’s estimate of theta (θw), Tajima’s D, and Fu & Li’s D* (11; 3). Tajima’s D test takes into account the average pairwise nucleotide diversity between sequences and the number of segregating sites expected under neutrality for a population at mutation-drift equilibrium (60). Tajima’s D is useful to detect departures from neutrality when considering an excess of rare alleles indicating positive selection/selective sweep, or the opposite, excess of common alleles that leads to assume BS has operated in the population. In our case, Tajima’s D helps to find polymorphisms at intermediate frequency. Watterson’s theta measures the population mutation rate which is understood as the product of the effective population size and the neutral mutation rate from the observed nucleotide diversity of a population (69). In this case, θw is an indicator of high level of polymorphisms. Fu & Li’s D* statistics considers the number of derived singleton mutations and the total number of derived nucleotide variants without an outgroup (19). We used a combination of these three test statistics to detect excess of common polymorphisms along the allele frequency spectrum relative to expectations under neutral equilibrium. The use of three indicators may seem overly conservative, but it helps to reduce false positives and to detect genes or genome regions that are robust candidates for operating under BS. Neutrality tests were calculated with VariScan 2.0.3 (30) using total number of segregating sites and excluding sites containing gaps or ambiguous nucleotides.

We performed a genome-wide scan to find genes or genome regions under BS using a sliding window approach. Thomas and colleagues (ref. 64) tested windows of two sizes, 100 bp and 200 bp for *S. aureus* genome analysis coming to the conclusion that 200 bp windows is the optimal and 100 bp windows is the second best alternative for a genome scan. The type strain of *S. aureus* subsp. *aureus* DSM 20231^T^ has a genome of 2,9 Mb (35) which is slightly smaller than the RSSC chromosome (3.7 Mb for reference strain GMI1000) (55). Moreover, the average length of protein-coding genes is similar for both bacterial species [946 bp (chromosome), 1,077 bp (megaplasmid) for RSSC and about 1,009 bp for *S. aureus;* see ref. 55 and 35]. Therefore, a 200 bp windows seems to be an adequate window size for RSSC. All three statistics were calculated for consecutive, non-overlapping, 200 bp windows, and only those windows with the highest 5% values coinciding in the three statistics were chosen as possible candidates for further analyses. Windows without single nucleotide polymorphisms (SNPs) among aligned genomes were excluded from analysis because the statistics are calculated based on polymorphisms.

Per site mutation (θ) and recombination (ρ) rates are parameters useful for understanding the recent history of RSSC populations, however they also help to test demographic models to discover which one best fits the observed data for each population (see below). These parameters were estimated using a penalized approximate likelihood coupled to a Bayesian reversible-jump Markov chain Monte Carlo sampling scheme. For this, we set up the starting ρ value to 30, penalized each block with a value of 10 and used the gene conversion model. We run 10^6^ chains to obtain ρ and θ values using the program INTERVAL (41) implemented in the RDP4 package (39). Because RDP was not designed to handle long genomic sequences, we estimated values of ρ and θ by averaging the obtained values from sets of 50,000 bp each along the length of nucleotide sequence alignments.

The summary statistics (θw, Tajima’s D, and Fu & Li’s D*) must be carefully analyzed because different demography scenarios could give similar signals as BS when applied to real population data. For example, different population structures like a contraction or a selective bottleneck could generate confounding indications mimicking BS. To correct potentially confounding effects of demography we need to select adequate null demographic models and test them with real data. For this purpose, we adopted a simulation-based approach to generate genetic statistics under three main demographic scenarios: standard neutral model (SNM), a recent population contraction model (PCM), and a recent bottleneck model (BNM). The SNM assumes a constant-sized population, thus Tajima’s D is expected to be zero (60). Under PCM and BNM assumptions, Tajima’s D is positive or shows higher values than with SNM, which indicates the abundance of prevalent lineages before a contraction or a bottleneck effect. Simulations were performed under the coalescent simulation framework by employing the algorithm described in Hudson (ref. 29) to infer the coalescent tree with recombination. The PCM assumes that the population has undergone a size reduction at a given time that we fixed at 0.005 coalescent time units before the present, according to Thomas and collaborators (ref. 64). Coalescent time units are measured in *4Ne* generations where *Ne* corresponds to the current effective population size (29). For BNM simulations, the model assumes that the population suffered two demographic events, a contraction and then a population growth. In this case, we calibrated time *(Tc* and *Tr* time of contraction and time of recovery, respectively) for first and second events as 0.005 coalescent time units before the present until a relevant demographic event (64). The reduction of population size (Ne) relative to constant growth was set to 5, for PCM and for the first and second demographic events of BNM. The fivefold reduction of the original population size is based on the *Ne* decrease reported in experimental studies performed on different bacterial species (47; 72; 10; 37). Finally, ρ and θ values calculated previously were used to complete the information required to run the simulations. For each window, we computed 10,000 coalescent simulations using DNASP v. 6.11.01 for the three summary statistics under the relevant demographic model (53). A p-value was estimated for each window to validate statistically the potential differences between simulated and observed data. Windows with extreme *(i.e.* significant) p-values (at the right tail, *p*_Sim<*obs*_ < 0.1 or *p*_Sim<*obs*_ < 0.05) for the three statistics and the three demographic models were recorded as highly significant and accepted as candidates under BS. However, windows with significance for only two statistics (Tajima’s D and Fu & Li’s D* or Tajima’s D and θ_w_, see Table 2 and Supplementary Table 3) were also accepted as secondarily significant.

### Gene identification and function

Sequences of windows with significant values were used to identify genes that overlap in them. For this, Blastn searches were performed using standard settings (2). We used four RSSC reference strains for sequence comparison and gene identifier assignation: GMI1000 for phylotype I; CFBP2957 for phylotype IIA; Po82 for phylotype IIB; and PSI07 for phylotype IV. Uniprot (63) and Pfam (17) databases including their tools were used to retrieve information on the features and function of proteins. The respective gene ontology (GO) term was applied to each identified protein using QuickGO (https://www.ebi.ac.uk/QuickGO/). The KEGG database was used for further understanding putative gene functions, utilities of the bacterial systems and to define orthologs for RSSC genes under BS (31). Identification of T3SS effector proteins was achieved using the web interface named “Ralstonia T3E” (https://iant.toulouse.inra.fr/T3E) with the curated effector repertoire database (45).

## RESULTS

### Genome sequence alignment and population parameters

For all analyses performed in this work, we chose to work with locally collinear blocks (LCBs) than with complete genome alignments because LCBs produce aligned and concatenated sequences composed of homologous regions of sequence shared by the genomes under study. In this way, only conserved segments that appear to be internally free from genome rearrangements were considered for population parameters and summary statistics calculations. This is critical for calculations aimed at detecting polymorphisms on aligned sequences. Genome alignments of the RSSC phylotypes analyzed in this work produced a variable number of LCBs that concatenated represent about (or higher than) 50% of their respective genomes (except for the megaplasmid of phylotype IV, see Table 1). Because some genome sequences from the database are poor in megaplasmid sequences, we were only able to align seven genome sequences for the phylotype IIA megaplasmid (Table 1).

**Table 1.** RSSC genomic sequences used in this analysis and population parameters and summary statistics calculated for whole replicons sequence data.

| Phylotype/<br>replicon | Number of<br>genomes<br>analyzed | Number of<br>nucleotides<br>analyzed | Percentage<br>of GMI1000<br>replicon<sup>a</sup> | $\rho$<br>(per<br>site)<sup>b</sup> | $\theta$<br>(per<br>site)<sup>b</sup> | $\rho/\theta$ | Tajima's<br>D | $\theta_w$<br>(per<br>site)<sup>c</sup> | Fu-Li's<br>D* |
| --- | --- | --- | --- | --- | --- | --- | --- | --- | --- |
| I/chromosome | 20 | 1907685 | 51.33 | 0.0120 | 0.0052 | 2.399 | -0.438 | 0.0051 | -0.676 |
| I/megaplasmid | 20 | 1282321 | 61.22 | 0.0233 | 0.0067 | 3.470 | -0.450 | 0.0067 | -0.743 |
| IIA/chromosome | 12 | 1971855 | 53.06 | 0.0008 | 0.0103 | 0.071 | 1.084 | 0.0101 | 0.895 |
| IIA/megaplasmid | 7 | 938400 | 44.80 | 0.0020 | 0.0164 | 0.125 | -0.642 | 0.0160 | -0.725 |
| IIB/chromosome | 20 | 1451109 | 39.05 | 0.0007 | 0.0092 | 0.071 | 0.923 | 0.0080 | 0.539 |
| IIB/megaplasmid | 20 | 927177 | 44.27 | 0.0010 | 0.0133 | 0.075 | 0.930 | 0.0116 | 0.522 |
| IV/chromosome | 5 | 1957952 | 52.68 | 0.0011 | 0.0112 | 0.088 | -0.434 | 0.0104 | -0.467 |
| IV/megaplasmid | 5 | 503195 | 24.02 | 0.0021 | 0.0139 | 0.136 | -0.420 | 0.0134 | -0.468 |

<sup>a</sup> As a reference, the GMI1000 chromosome has 3,716,413 bp and the megaplasmid 2,094,509 bp (ref. 55).
<sup>b</sup> $\rho$ and $\theta$ : per site recombination and mutation rate, respectively.
<sup>c</sup> $\theta_w$ : Watterson's estimate of theta

Alignments were analyzed for information on population parameters which are necessary for the simulations (see below). Per site recombination rate (ρ) and per site mutation rate (θ) vary across different phylogenetic groups in RSSC (Table 1). The chromosome of phylotype IIB and the chromosome of phylotype I show the lowest value for ρ and θ respectively. On the other hand, the highest values of the two parameters are shared by the megaplasmid of phylotype I (for ρ) and the megaplasmid of phylotype IIA (for θ). Interestingly, the relation ρ/θ gives opposite values depending on the phylotype. Phylotypes II (A and B) and IV show values lower than 1 for ρ/θ, while phylotype I reaches values higher than that. This result suggests that the role played by recombination seems to be uneven across RSSC lineages and that recombination had a stronger influence on introducing nucleotide substitution relative to mutation in phylotype I (both replicons) than in other phylotypes.

### Summary statistics

RSSC genome alignments were scanned for BS signatures in both replicons *(i.e.* chromosome and megaplasmid). We focused the analysis on phylotype I, II and IV as there were not enough genome sequences available in the databases for phylotype III at the time of the analysis and phylotype II was analyzed in both its subclusters as they were separate and independent phylogenetic groups (Table 1). The extent of polymorphism was measured by using the three summary statistics mentioned above (θw, Tajima’s D, and Fu & Li’s D*). The Tajima’s D values calculated for the whole replicon of each phylotype ranged from −0,6417465 to 1,084 depending on phylotype (Table 1). Phylotypes I and IV show Tajima’s D distribution shifted towards negative values in both replicons, as well as phylotype IIA (megaplasmid). On the contrary, phylotypes IIA (chromosome) and IIB (both replicons) show a tendency towards positive values. Fu & Li’s D* results follow a similar pattern as Tajima’s D. This suggests that both these statistics are highly correlated, an aspect that is confirmed later (see below). When we estimated the summary statistics using the sliding window strategy, an ample assortment of values was obtained for each phylotype and replicon. After eliminating windows without SNPs, we observed extreme Tajima’s D values (such as 3.46 and −2.506 for the chromosome in phylotype I) but also moderate values, along all windows analyzed (Supplementary Table 1). The tendency towards negative values was reflected in Tajima’s D and Fu & Li’s D* mean values of sliding windows analysis for phylotypes I (both replicons), IIA (megaplasmid) and IV (both replicons) (Supplementary Table 1, Figure 1). Watterson’s θ values are relatively high for all phylotypes except for phylotype I and IIB (chromosome). The slight differences between θ and θw observed in Table 1 are due to the way of calculating this statistic, as in one case, we employed the Bayesian method and in the other the formula proposed by Watterson (ref. 69).

**Figure 1.**
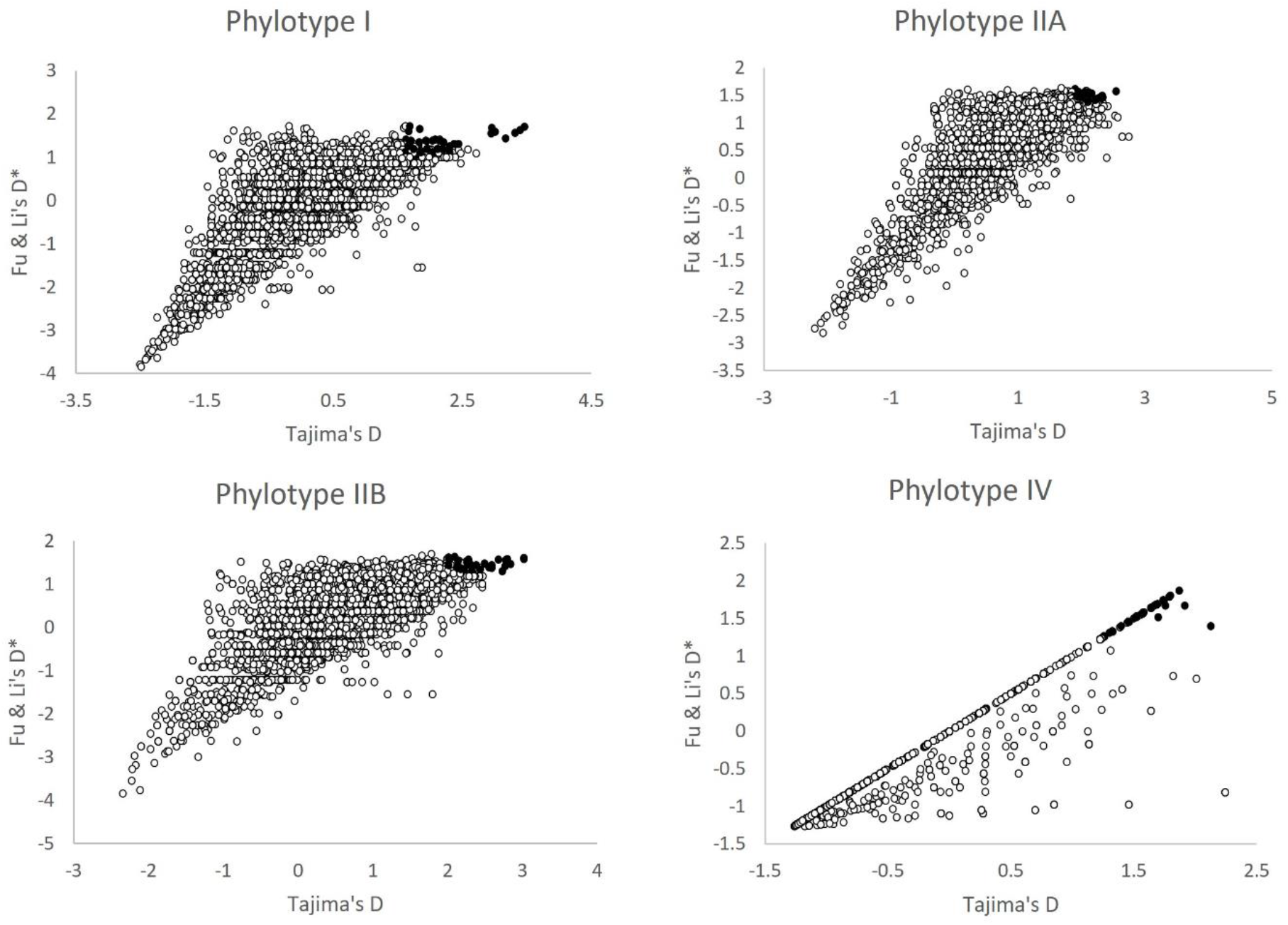
Two-dimensional plot of Tajima’s D and Fu & Li’s D* values for all windows. Shaded dots represent the 5% top windows of the distribution of both measures of BS.

A two-dimensional plot of all three statistics suggests that their values are correlated (Figure 1). To confirm a possible correlation between them, we calculated the Spearman rank correlation coefficient between θw, Tajima’s D and Fu & Li’s D* using the sliding window data. As expected, results show that there is a strong pairwise correlation among the three statistics for all phylotypes and replicons except for phylotype I when comparing θw and Tajima’s D (Supplementary Table 2). In some cases, a very high positive correlation was observed, as is the case of phylotypes IIA and IV for Tajima’s D-Fu & Li’s D* combination (0.738, 0.964 and 0.982, 0.976, respectively) suggesting a strong agreement between these statistics. This result also supports the idea that the high values of the statistics point out to real BS signatures (or demographic structuring) in aligned sequences rather than being random values.

### Simulations and candidate genes under balancing selection

We tested whether the unusual incidence of high values of summary statistics obtained from aligned sequences was due to BS on RSSC genomes or reflected effects of demography. We adopted the widely used strategy based on simulation of genetic data under the coalescent framework. Three most plausible demographic scenarios were tested (SNM, PCM and BNM) as null models. Although these models may not represent the exact history of RSSC populations because of their intrinsic complexity, this approximation is sufficiently advantageous to be used as a null demographic model focused upon reducing false positives. We included in our analysis the gene cluster *agr* from *S. aureus* as a positive control (64). We analyzed 3,537 bp of the *agr* cluster using the standard procedure for BS signature detection in RSSC aligned sequences as detailed in the Data and Methods section. This analysis produced 18 windows, however, in none of them, we obtained maximum matching values for the three statistics. As expected, windows with high observed values of Tajima’s D, θw, or Fu & Li’s D* showed very significant values after simulations according to the demographic models tested in this work (observed values: Tajima’s D= 2.72677**; θ_w_= 0.05249**; Fu & Li’s D*= 1.73125**, the double asterisk meaning significant difference at p<0.05 compared to values obtained with simulations for SNM). After having demonstrated confidence in the analysis using this positive control, we applied the same procedure to scan the RSSC aligned sequences. Results show (Table 1) that the top 5% of the distribution of the summary statistics exceed the respective simulated values (under the corresponding demographic model) in most of the cases, as validated by hypothesis testing significance. Note that the power of this detection resides in the concurrent consideration of all three statistics, Tajima’s D, θw, and Fu & Li’s D*. This result provides a robust evidence that the windows with high values of summary statistics correspond to genes or genomic regions under BS (Table 1). Subsequently, we identified the genes overlapping the candidate windows (Table 2). A list of unidentified genes (unknown gene function) or windows corresponding to intergenic regions is detailed in Supplementary Table 3.

**Table 2.**
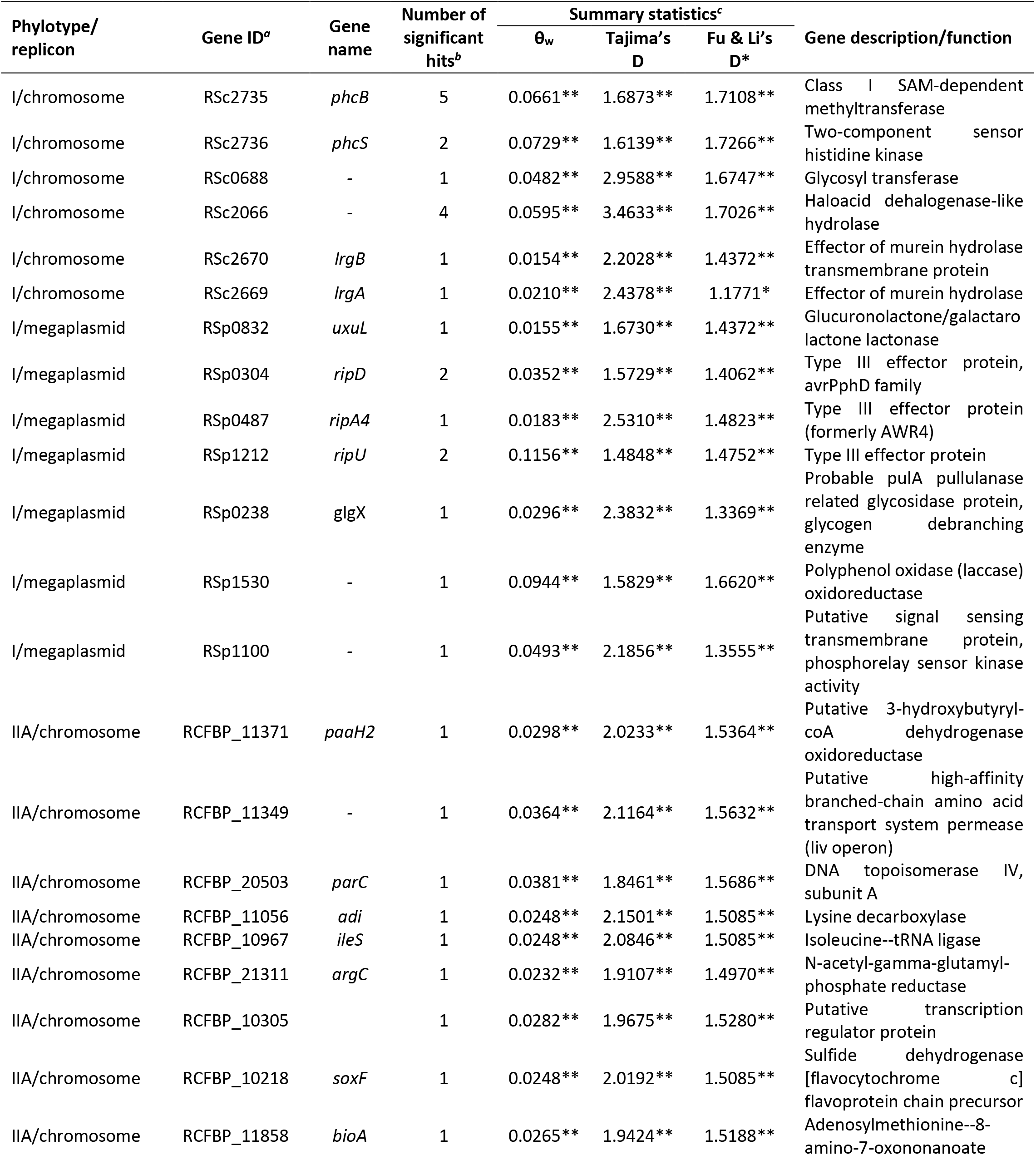
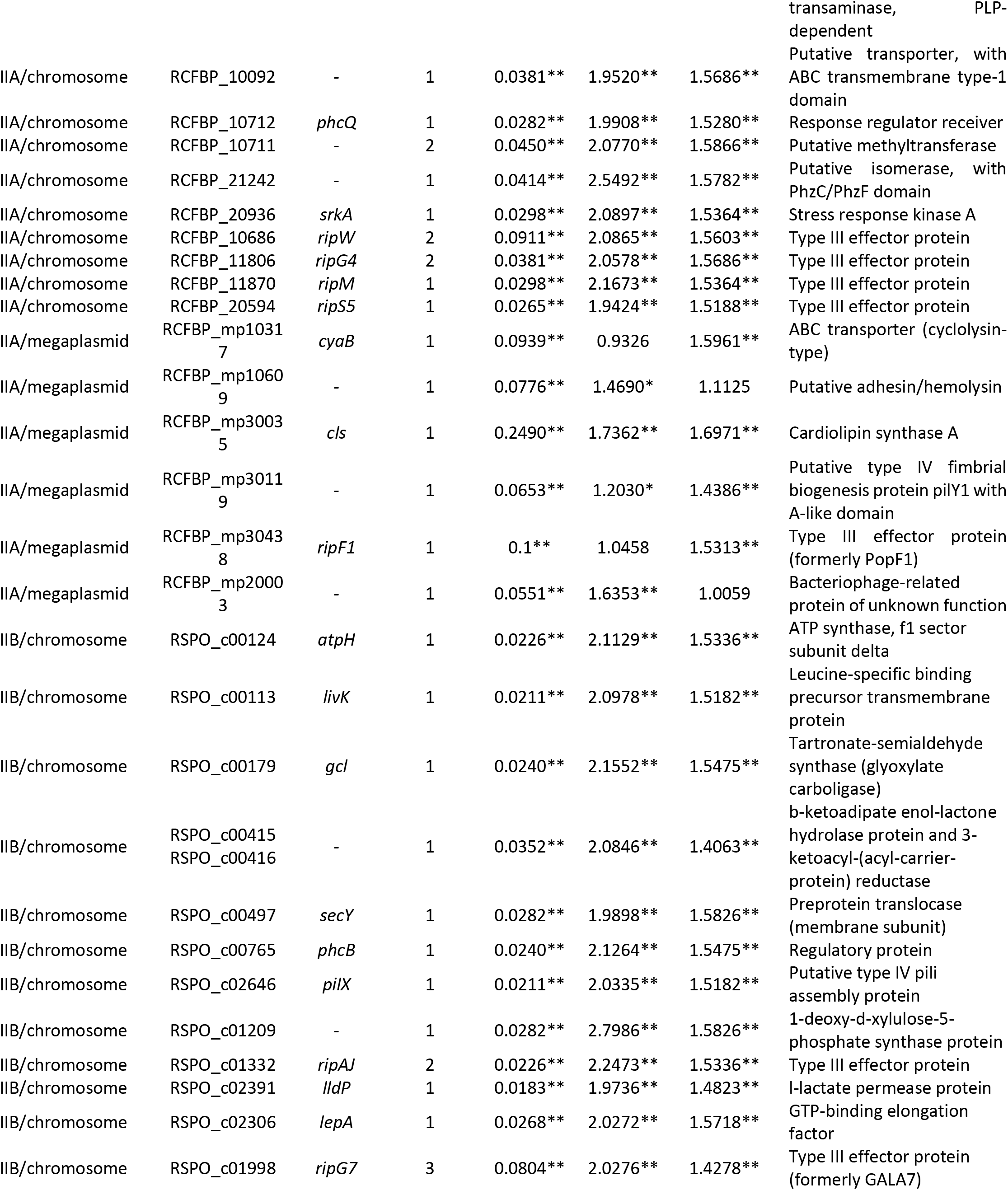
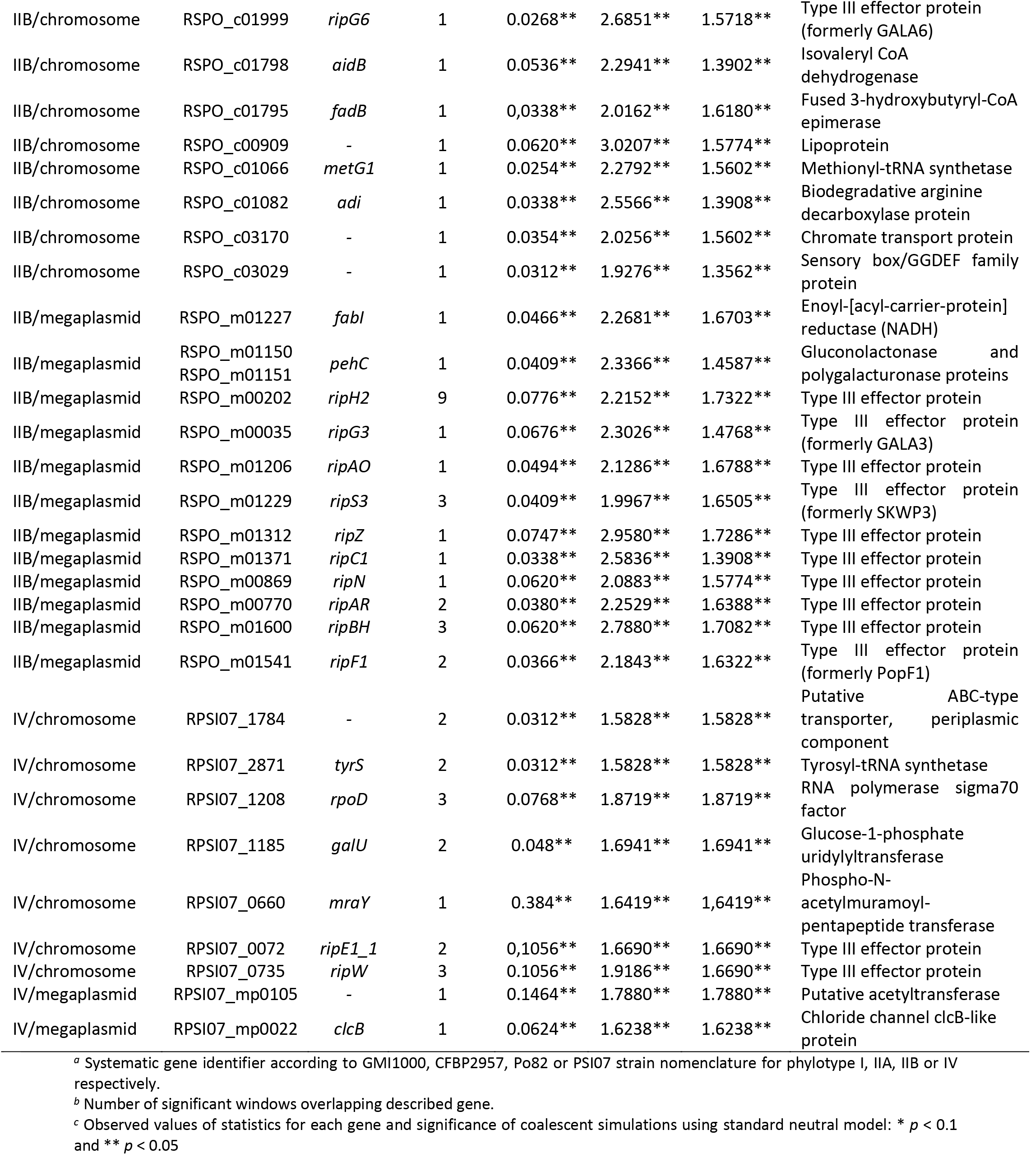
Identity and probable function of genes showing highest observed values of three statistics (θw, Tajima’s D, and Fu & Li’s D*) in the genome-wide analysis of RSSC phylotypes.

In general terms, the results show that BS affects more frequently coding regions than non-coding sequences in RSSC genomes (compare Table 2 with Supplementary Table 3). We found 161 windows with significant values for the three statistics. Demography simulations reduced the number to 142 significant windows that correspond to 78 known genes (Table 1) and 22 intergenic regions or genes with unknown identity or function (Supplementary Table 3). This may indicate that 19 windows are probably false positives. The percentage of genes detected under BS is low (1.7%, 78 genes out of 4,585 which is the median protein-coding genes in RSSC according to the Genome Database, https://www.ncbi.nlm.nih.gov/genome/?term=Ralstonia+solanacearum). This result is consistent with other analyses in eukaryotic systems like humans (4) or plants (52) and also in prokaryotes (64) that stress the rarity of finding BS signatures on sequence genomes. The candidate genes under BS are described below according to phylotype and replicon.

*Phylotype I.* We detected 440 windows for the chromosome of this phylogenetic group at the top 5% of the distribution, however only 21 showed concurrent high values in all three summary statistics and 14 were recorded as highly significant after the simulation process.

We found seven peak values of Tajima’s D, θw and Fu & Li’s D statistics on *phcB* (five extreme values) and *phcS* (two extreme values, Figure 2, Table 2) genes. These two genes are arranged in an operon together with a third gene named *phcR.* The gene *phcB* encodes a SAM-dependent methyltransferase that synthesizes 3-hydroxy palmitic acid methyl ester (3-OH PAME), a quorum-sensing signal that accumulates in the extracellular space when the bacteria are multiplying rapidly in a restricted space (18). Quorum-sensing is a key process regulating and synchronizing the expression of specific genes involved in biofilm formation, pathogenicity, and production of secondary metabolites like siderophores, exoproteases, and exotoxins (36). *PhcS* (histidine kinase) and *phcR* (response regulator) genes code for elements of a two-component regulatory system that responds to threshold concentrations of 3-OH PAME by elevating the level of functional PhcA, the fourth component of the system (8). PhcA is the global virulence regulator in RSSC since it regulates hundreds of genes directly involved in pathogenesis but also in basal metabolism and cell homeostasis (34; 48).

**Figure 2.**
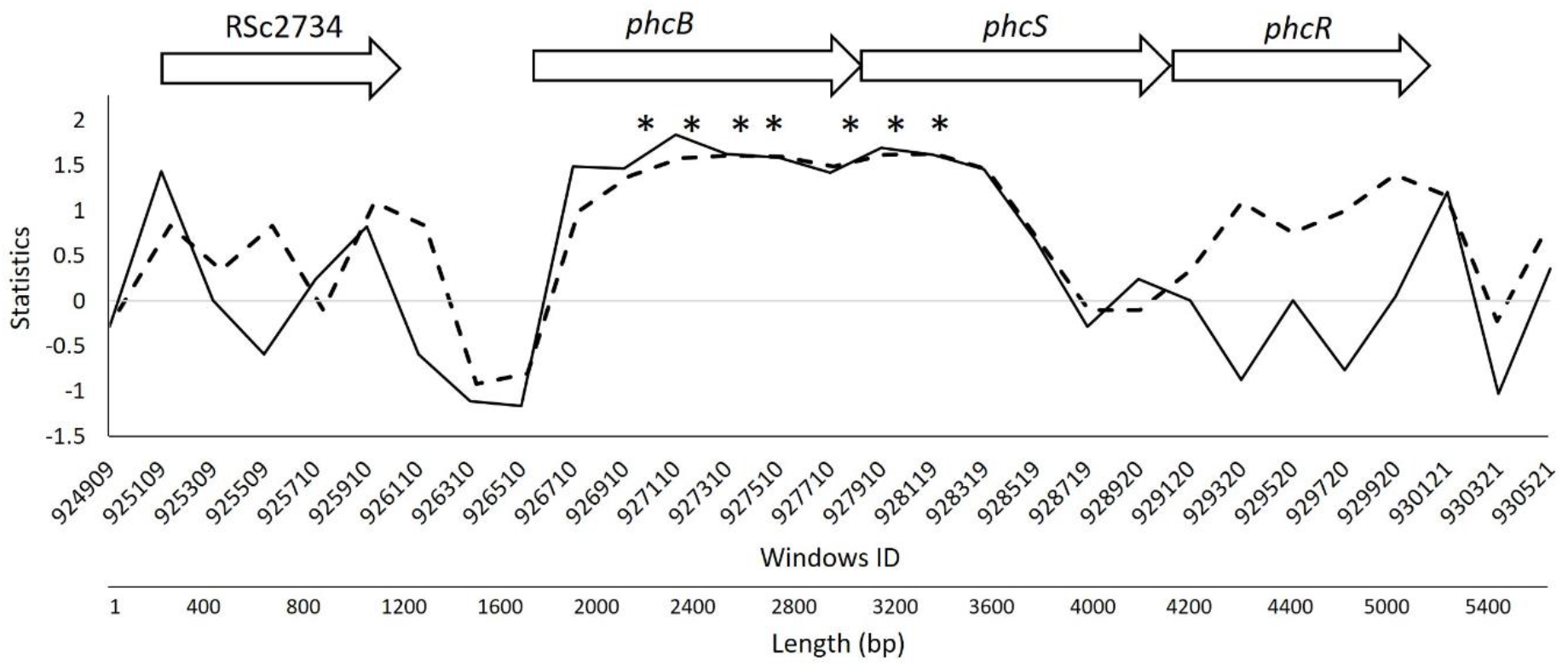
Analysis of genomic region corresponding to the *phcBSR* operon in strain GMI1000 showing sliding window analyses for two statistics: Tajima’s D (solid line) and Fu & Li’s D* (dotted line). All three genes of the operon comprise about 3.9 Kb of the genome; however, the gene (RSc2734) physically preceding this operon is also shown for comparison purposes. Windows are 200 bp and asterisks indicate the windows with extreme values of respective statistics. Arrows represent gene arrangement in the genome of strain GMI1000.

Another gene showing multiple peaks in statistics values is RSc2066 that codes for a haloacid dehalogenase-like hydrolase (Table 2). In this case, four consecutive high values of the statistics suggest that this gene is likely under BS. This enzyme has a hydrolase activity that cleaves different bonds (i.e. C O, C-N, C-C), however its exact role at the cellular level is unknown.

In this phylotype we also found genes related to basic metabolism like a glycosyl transferase and an operon consisting of two genes, *lrgAB,* that modulates murein hydrolase activity which is linked to biofilm dispersal and cell lysis (24). These *lrgAB* genes intervene indirectly in pathogenesis since an essential step in this process is the formation and dispersal of biofilms in RSSC (36).

For the megaplasmid we found 304 out of 6162 windows with highest Tajimas’s D, θw, and Fu & Li’s D* values. After simulation for relevant demographic models, only nine windows generated significant values.

Some interesting genes associated to virulence were observed in this replicon (Table 2). We identified three different T3SS effector genes as targets for BS: *ripD* of the avrPphD family; *ripA4* and *ripU.* Interestingly, both *ripD* and *ripU* show two significant hits (two windows with significant values) along their coding sequences. *RipU* is part of the core-effectome within the RSSC as well as *ripA4* that is common in effector collections and plays an important role in the interaction between *R. solanacearum* and the pepper plant (46). Another gene, *uxuL* (RSp0832) codes for the main glucuronolactone/galactarolactone lactonase in the genome of the GMI1000 strain. UxuL is organized within an operon with three other genes: *garD* encodes a D-galactarate dehydratase, RSc0831 a putative NAD-dependent epimerase/dehydratase and *pehC* a polygalacturonase. PehC is an enzyme related to virulence since it cleaves oligomers of galacturonate, however its exact role is unknown. It was hypothesized that PehC acts by degrading plant oligogalacturonate signal molecules that elicit production of reactive oxygen species (ROS) as a defense response. This degradation would reduce tomato antimicrobial responses and increase bacterial virulence (23). This operon is regulated by GulR, a transcription factor of the LysR family involved in glucuronate utilization and metabolism. Downstream of this operon is located *exuT,* the galacturonate transporter gene.

Conversely, genes that are not directly related to virulence but to primary metabolism were also identified in megaplasmid aligned sequences: A probable pullulanase related glycosidase protein (PulA) that might work like a glycogen debranching enzyme; a polyphenol oxidase (laccase) oxidoreductase and a putative signal sensing transmembrane protein with phosphorelay sensor kinase activity. Lastly, a significant window matched with an intergenic region surrounded by a hybrid sensor histidine kinase/response regulator and upstream of an integrase related to phage or transposon insertion (Supplementary Table 3).

*Phylotype IIA.* At the chromosome level we selected 444 windows that showed 5% highest scores in each statistic. From these, 21 windows showed highest values for all three statistics and also significant values on simulations with respective demographic models.

The first genes that appear in the list are those involved in essential cell functions. There are various enzymatic functions *(i.e.* a 3-hydroxybutyryl-coA dehydrogenase oxidoreductase, an isoleucine-tRNA ligase, a transcription regulator and others, Table 2) and diverse transporters (a permease from the *liv* operon, a binding-protein-dependent transporter). Among this group, a gene that attracted our attention is *adi* which encodes a lysine decarboxylase (LDC). This gene and other related genes (arginine and ornithine decarboxylases) are directly involved in amino acids metabolism but indirectly in pathogenesis. Studies on other bacterial species indicate that these genes are implicated in stress response against the low pH in the medium (44; 43) and against oxidative stress and chemical quenching induced by the host (66). This gene product or LDC metabolic products also intervene in cell adhesion to host tissues (67)

Among the genes related to virulence and survival, we found two contiguous genes, *phcQ* and another one downstream from it showing elevated values of selection statistics. PhcQ is a response regulator receiver, from the CheY family and part of the *phcBSRQ* operon that regulates PhcA, the master regulator that positively and negatively regulates many genes responsible for pathogenicity in RSSC (70). The gene contiguous to *phcQ* encodes a methyltransferase, however it is not known if PhcQ participates in quorum sensing as does the main methyltransferase, PhcB. Two additional genes were associated to BS signatures: *srkA* and RCFBP_21242. The *srkA* gene encodes a stress response kinase A, which probably counteracts the accumulation of ROSs produced by the host and protects the bacterial cell from antimicrobial and environmental stressors in a similar way to the YihE protein kinase of *Escherichia coli* (14). RCFBP_21242 encodes a putative isomerase with a phenazine biosynthesis (PhzC/PhzF) domain. Phenazines constitute a large group of nitrogen-containing heterocyclic compounds produced by bacteria and show an ability to handle ROS, contribute to biofilm formation, cell adhesion and enhance bacterial survival, among other activities (50).

Results on the T3SS effector repertoire analysis of phylotype IIA-chromosome showed a number of genes with a BS signature: *ripM, ripW, ripG4* (formerly GALA4) and *ripS5* (formerly SKWP5). *RipG4* and *ripW* were associated to two significant windows each suggesting these genes are clearly under BS. Since we have used the CFBP2957 strain as reference for gene identification, we find that this strain has an insertion of a transposon encoding a transposase (RCFBP_20595) in the *ripS5* gene, therefore this appears to be a pseudogene copy of this effector. Most of the phylotype IIA strains show a disruption in the *ripS5* gene due to transposon insertions, however there are some strains harboring the complete gene *(i.e.* the RS_489 strain)(45).

At the megaplasmid level, phylotype IIA showed six genes with significant signatures of BS after filtering with coalescent simulations: one related to basic metabolism *(cyaB,* an ABC transporter) and four pertaining to pathogenicity: a putative adhesin/hemolysin that plays a significant role in cell adhesion; a cardiolipin synthase A, from the phospholipase D family, involved in membrane biosynthesis and toxin production and resistance (61); a putative Type IV fimbrial component, encoded by the *pilY1* gene participating in Type IV pili biosynthesis (type IV pili are essential for adhesion and pathogenesis)(1), and a bacteriophage-related protein with unknown function. Finally, a T3SS effector named *ripF1* [formerly PopF1] that is very well characterized (42).

*Phylotype IIB.* Three hundred fifty two windows corresponding to the top 5% of the distribution were analyzed for the chromosome. Only 33 windows showed highest values of the three statistics concurrently, but 23 windows showed significant values after coalescent simulations.

The most abundant group of genes identified in this chromosome are those implicated in primary metabolism with an ample diversity that varies from genes encoding metabolic enzymes (synthases, epimerases, etc.) to a number of permeases and other transporter related genes (Table 2). Again, an amino acid decarboxylase was found within this group.

Various genes are linked to virulence: a key component of pili biogenesis (Type IV pili assembly protein PilX) and the gene responsible for the production of the molecule that mediates quorum sensing, *phcB* were identified. Among genes encoding T3SS effectors, three were most notable *(ripAJ, ripG6* and *ripG7)* and multiple windows enriched two of them (two and three hits for *ripAJ* and *ripG7,* respectively, Table 2).

Interestingly, a conserved protein (RSPO_c02827) showed also two significant hits along its sequence but its function is unknown (Supplementary Table 3).

We identified 26 windows with significant values distributed across the megaplasmid after the simulation process. Since many virulence-related genes reside in the megaplasmid, it was not surprising to have identified many of them. Ten different T3SS effector genes were found (Table 2) and some were noted by redundant windows as is the case of genes *ripH2* (9 hits), *ripS3* (3 hits), *ripBH* (3 hits), *ripAR* (2 hits) and *ripF1* (2 hits). On the other hand, only few genes involved in basic metabolism were identified: an enoyl reductase (NADH dependent) and two contiguous genes, polygalacturonase and gluconolactonase, that overlap within a single window (N-terminus of the first and C-terminus of the second enzyme).

*Phylotype 4.* The chromosome showed 463 windows in the top 5% of the distribution for each statistic, and after selection for the matching values in the three statistics and the simulation, only 15 were retained as highly significant for further analyses.

We found interesting genes in the chromosome such as one encoding the RNA polymerase sigma 70 factor which gathered three consecutive windows. Other genes that received multiple hits include a tyrosyl-tRNA synthetase, a glucose-1-phosphate uridylyltransferase and a putative ABC-type transporter. On the other hand, a phospho-N-acetylmuramoyl-pentapeptide transferase was detected by one window. In the gene group related to virulence, we found two T3SS effector genes with multiple windows: *ripE1* from the AvrPphE family and *ripW* [formerly PopW], a hairpin with a pectate lyase domain.

At the megaplasmid level, we found only two metabolically essential genes with significant values: a putative acetyltransferase and a chloride channel clcB-like protein.

## DISCUSSION

In this work, we report the systematic exploration of the genomes belonging to the main RSSC phylotypes with the intention of finding signatures of BS. To our knowledge this is the first time that a bacterial plant pathogen is analyzed for this type of selection at the genomic level. The analysis was performed on the main replicons of RSSC (chromosome and megaplasmid), but not on small plasmids, phages or mobile genetic elements. We scanned genome sequences using a sliding window approach and subsequently applied widely used summary statistical tests aimed at detecting the excess of polymorphisms on 200 bp-window sequences: Watterson estimator theta, Tajima’s D, and Fu & Li’s D*. We chose to use these tests rather than other strategies (i.e. model based methods) because of their simplicity, wide range of BS forms detected and broad access to diverse software tools. This strategy together with exhaustive coalescent simulations to correct confounding effects of demography was an effective approach to reach our objective to detect genes and genomic regions under BS in RSSC. Tajima’s D is useful for detecting intermediate and ancient signatures of BS. In contrast Fu & Li’s D* and θw help to identify relatively recent instances of this type of selection. Our approach may be overly conservative, and hence we might have missed some genuine occurrences of BS. However, it may have conferred more certainty to the positive hits found on RSCC genomes. Indeed, we detected dozens of gene candidates in RSSC genomes in agreement with Fijarczyk and Babik (ref. 16) who recognized this is common in pathogens’ genomes.

The results confirm the validity of the methodological strategy used and add new insights to understand RSSC and plant host interaction. We have found many bacterial genes that show unambiguously features of being under BS. The *phcBRS* operon scored 7 significant windows in phylotype I and one in phylotype IIA and one in phylotype IIB, indicating this genomic region is under strong BS. Remarkably, Guidot and collaborators (ref. 25) also found that one component of this system, *phcS,* was subject to strong selection from the plant host given the evidence that this gene was targeted by mutations in an *in planta* experimental evolution system. The *phcBRS* is responsible for controlling a complex regulatory network that responds to environmental conditions and bacterial cell density. This system comprises a two-component signaling system composed of a histidine kinase, PhcS, that phosphorylates the response regulator, PhcR, when the signal molecule has reached the threshold level and PhcB is the enzyme responsible for forming the signaling molecule that mediates a quorum sensing communication. The main quorum sensing signaling molecules are 3-OH PAME (methyl 3-hydroxypalmitate) or 3-OH MAME (methyl 3-hydroxymyristate) depending on the producer strain (28). The key player in this network is a global transcriptional regulator, PhcA, that coordinates the expression of several virulence-related genes including those responsible for the major extracellular polysaccharide, cell wall degrading enzymes, T3SS effectors, and others representing a total of 383 genes (48). Interestingly, an equivalent but simpler network in *S. aureus,* the *agr* locus, is also a two-component signal transduction system (membrane-bound histidine kinase sensor, AgrC and transcriptional regulator, AgrA), with a signal molecule (an auto-inducing peptide, AgrD) and a protein responsible for the maturation and export of the signal molecule (AgrB). Again, the key component in this system is the master transcriptional regulator AgrA that binds onto the promoter region and induces transcription from two divergent promoters, P1 and P2 (65). Although this system does not show homology at the sequence level with the *phcBRS* system in RSSC, it is functionally analogous since it leads to up and down-regulation of over 70 genes, 23 of which are known to be directly related to virulence (21). Interestingly, the *agr* locus has the strongest known signatures of BS in bacteria to date due to the high number of common polymorphisms. For this reason, the *agr* locus has been proposed as the positive control of BS for further studies in bacteria (64).

We have also found a set of genes with strong BS signatures whose function is related to adhesion, motility and biofilm formation. Genes encoding Type IV fimbrial biogenesis proteins were found in phylotype IIA (megaplasmid, *pilY1)* and phylotype IIB (chromosome, *pilX).* These proteins are essential for the assembly and function of Type IV pili, filamentous structures that mediate bacterial adhesion to surfaces including host cells. This adhesion is tightly linked to the bacterial pathogens’ ability to promote the formation of microcolonies and biofilms as well as to their twitching motility and virulence (32; 57). In phylotype I (chromosome), we found two LytSR-regulated genes, *lrgA* and *lrgB* that code for a murein hydrolase exporter and a protein having murein hydrolase activity respectively (24). Both proteins are required for biofilm dispersal that is accompanied by cell lysis and death. Biofilm formation and disruption is a critical step in the process of infection and pathogenesis for RSSC strains. Diverse types of molecules mediate the release of the cells from biofilms, including degrading enzymes (among them, murein hydrolases), nucleases and others (40; 68). Additionally, we identified one gene under BS that seems to be directly related to the biosynthesis of phenazines in phylotype IIA. Phenazines constitute a large group of nitrogen-containing heterocyclic compounds produced by a wide range of bacteria, with diverse physiological functions. Among these, they influence swarming motility and biofilm architecture through a not fully understood mechanism (51).

T3SS effectors are key virulence factors at the forefront of the arsenal that RSSC strains harbor to infect plants and achieve full pathogenicity including the metabolic adaptation to parasitic life in the plant (9). T3SS effectors are delivered to plant cells through a proteinaceous needle-like structure, and once inside, they manipulate plant cell metabolism to suppress or evade defense responses and promote bacterial multiplication (6). *R. solanacearum* strains possess a large repertoire, with 94 effectors identified among RSSC sequenced (45). We found an ample collection of T3SS effector genes with moderate to very strong BS signatures in all phylotypes studied here (Table 2). Some of them belong to very well-known families of effectors like the GALA *(ripG4,* in phylotype IIA, chromosome; *ripG6* and *ripG7* in phylotype IIB, chromosome; *ripG3,* in phylotype IIB, megaplasmid), SKWP *(ripS5* in phylotype IIA, chromosome; *ripS3,* in phylotype IIB, megaplasmid), HLK (ripH2 in phylotype IIB, megaplasmid) and PopF type III translocators (ripF1). Interestingly, some effectors overlap with more than two windows and in different phylotypes *(ripW,* in phylotype IIA, chromosome and phylotype IV, chromosome), or in the same phylotype *(ripD*, phylotype I, megaplasmid *ripU,* phylotype I, megaplasmid, *ripW* and *ripG4* in phylotype IIA, chromosome; *ripAJ* and *ripG7* phylotype IIB, chromosome; *ripH2, ripS3, ripAR, ripBH* and *ripF1* in phylotype IIB, megaplasmid; *ripE1_1* and *ripW* in phylotype IV, chromosome), providing strong evidence that these genes are under BS.

Although genes dedicated to tasks of basal metabolism may seem less relevant for pathogenesis, they also play an important role in the interaction with the plant host and virulence. Peyraud and collaborators (ref. 49) developed a model system to study robustness and metabolic responses to internal and environmental perturbations in *R. solanacearum.* One of their findings highlight the active participation of primary metabolism in sustaining virulence, by activating functionally redundant reactions which may require redundant alleles to satisfy cellular demands including virulence. The expression of virulence factors (such as the exopolysaccharide) is controlled by the virulence regulatory network (VRN) that operate overlapping genes or operons involved in amino acid synthesis (49). While we did not particularly seek redundant or duplicate alleles in this work, we found a number of genes of primary metabolism that perform similar functions at the cellular level. For example, in the set of genes showing BS signatures there are two glucuronolactonases (carbohydrate metabolism), two aminoacyl-tRNA synthetases and two aminoacyl-decarboxylases (amino acid metabolism). These genes have roles in primary metabolism and probably are indirectly playing an essential role in virulence. Another group of genes that we should not neglect are those involved on defense and reduction of toxicity by metabolites produced by the plant host defense mechanisms. In the list of candidate genes under BS we can count a stress response kinase A *(srkA)* and a number of membrane transporters (ABC transporters and other permeases, see Table 2). Genes participating on defense pathways were also enriched in *S. aureus* genome analysis for BS signatures (64).

Analyses of BS operating on bacterial pathogens are particularly relevant to understand the dynamics of plant-microbe interactions. Host-pathogen coevolution leads to maintenance of high variation in genetic selected sites and nearby sequences. In plants, some loci involved in defense processes have highly polymorphic sequences that favor the occurrence of different resistance alleles (7). In pathogens, there is an equivalent scenario in which the pathogen maintains a high variation of polymorphisms in order to take advantage of the plant; consequently, the host-pathogen coevolution directs towards stable balanced polymorphisms and a high number of alleles in both host and pathogen populations (known as the trench warfare model)(59). Interestingly, in RSSC genomes, we found high variation in T3SS effector genes and other virulence-related genes as measured by Tajima’s D and other complementary statistics (Table 2), which may be under significant selection pressure by the plant host. Considering that RSSC has the ability to infect a large number of different plant species (20), it is not rare to find this high variation in the virulence factors. Some effectors (the so-called avirulence proteins) are recognized by proteins encoded by the plant R genes, however escape from host recognition is possible through fixing mutations on genes coding for effectors or other virulence proteins that increase variation. In order to evade plant detection and defense response, RSSC may tend to favor the maintenance of various allele alternatives (observed in the form of BS), which at the same time increases pathogen fitness. In a more applied sense, the identification of genes under BS, as illustrated in this work, opens the possibility to develop strategies towards establishing long term resistance or tolerance to pathogens in plants. These genes are potential targets for plant immunity, hence potential candidates to engineer broad disease resistance in agriculturally relevant plants.

## Acknowledgement

We thank Mrs. Helen M. Guigues for her valuable assistance with tables and figures.

## Funding Statement

The authors received no specific funding for this work.

